# Graph Convolutional Networks for Epigenetic State Prediction Using Both Sequence and 3D Genome Data

**DOI:** 10.1101/840173

**Authors:** Jack Lanchantin, Yanjun Qi

**Affiliations:** University of Virginia, Charlottesville VA 22903, USA {, }

**Keywords:** Genomics, Hi-C, Deep Learning, Graph Convolutional Networks

## Abstract

Predictive models of DNA epigenetic state such as transcription factor binding are essential for understanding regulatory processes and developing gene therapies. It is known that the 3D genome, or spatial structure of DNA, is highly influential in the epigenetic state. Deep neural networks have achieved state of the art performance on epigenetic state prediction by using short windows of DNA sequences independently. These methods, however, ignore the long-range dependencies when predicting the epigenetic states because modeling the 3D genome is challenging. In this work, we introduce ChromeGCN, a graph convolutional network for epigenetic state prediction by fusing both local sequence and long-range 3D genome information. By incorporating the 3D genome, we relax the i.i.d. assumption of local windows for a better representation of DNA. ChromeGCN explicitly incorporates known long-range interactions into the modeling, allowing us to identify and interpret those important long-range dependencies in influencing epigenetic states. We show experimentally that by fusing sequential and 3D genome data using ChromeGCN, we get a significant improvement over the state-of-the-art deep learning methods as indicated by three metrics. Importantly, we show that ChromeGCN is particularly useful for identifying epigenetic effects in those DNA windows that have a high degree of interactions with other DNA windows.

## 1 Introduction

The human genome includes over 3 billion base pairs (bp), each being described as A,C,G, or T. Chromatin (DNA and its organizing proteins) is responsible for many regulatory processes such as controlling the expression of a certain gene. Active chromatin elements such as transcription factors proteins binding at particular location in DNA or histone modifications are what constitute that location’s “epigenetic state”^1^. Understanding the epigenetic state of a local region of DNA is a step toward understanding how that region influences relevant gene regulation since epigenetic state is a direct factor in regulating expression. Since biological experiments are time-consuming and expensive, computational methods that can accurately simulate and predict the epigenetic state are crucial. Modeling the epigenetic state of each base pair has been a long standing challenge due to the sheer length and complexity of genome DNA. Deep neural networks have shown state-of-the-art performance in extracting useful features from segments of DNA to predict the epigenetic state (e.g., if a transcription factor protein binds to that location or not) [36, 1]. However, these methods heuristically divide DNA into local “windows” (e.g., about 1000bp long) and predict the states of each window independently, disregarding the effects of distant windows. Due to the spatial 3D organization of chromatin in a cell, distal DNA elements (potentially over 1 million bp away) have shown to have effects on epigenetic states [23, 21].

Fig. 1(a) illustrates the importance of using both sequence and 3D genome data. This figure shows long-range dependencies between chromatin windows, where the colored shapes represent multiple transcription factors (TF) proteins. TFs typically bind to specific sequence patterns in DNA known as motifs [29]. However, a TF may also bind to a DNA window due to the presence of other TFs nearby in the 3D space because they form a protein complex [5, 21]. Such a case will result in motifs far away in the 1D genome coordinate space, but nearby in the 3D space. The corresponding dependencies between the chromatin windows are illustrated by the triangle, square, and circle TFs. The two segments interacting in the middle of the diagram are very far in the 1D sequence representation (represented by the grey line), but very close in the 3D representation. Similarly, a TF may not bind to a segment with its motif present due to another interfering TF nearby in the 3D space. These types of interactions are lost in data-driven prediction models that only consider local DNA segments independently.

**Fig. 1:**
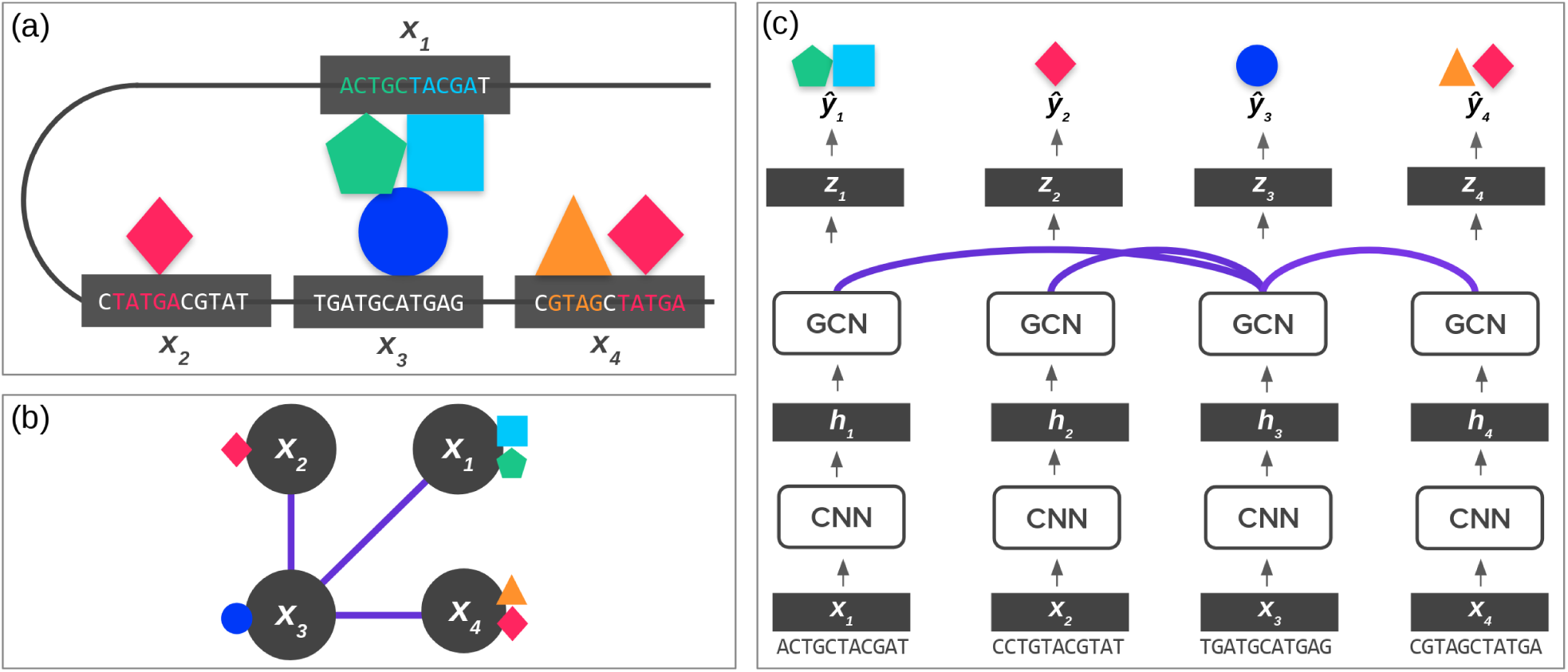
**(a) 3D Genome.** The 3D shape of chromatin can lead DNA “windows” (shown in grey boxes) far apart in the 1D genome space to be spatially close. These spatial interactions can influence epigenetic states, such as TFs binding (as shown by the colored shapes). In most cases, the DNA sequence determines the epigenetic state. However, it can also be influenced by interactions, such as the formation of TF complexes shown in the middle. **(b) Graph Representation of DNA.** Using Hi-C data, we can represent subfigure (a) using a graph, where the lines between windows are the edges indicated by Hi-C data. **(c) ChromeGCN.** By using a graph convolutional network on top of convolutional outputs the model considers the known dependencies between long-range DNA windows. The lines between windows correspond to edges in Hi-C data.

However, modeling these known long-range interactions between windows is difficult. Local sequence window-based prediction methods assume data samples are independent according to the commonly used independent and identically distributed (IID) assumption. Yet, the long-range dependencies existing in DNA make windows not IID.

Modeling long-range, or non-local interactions, has had a long history in many areas such as natural language processing, where the label of one particular segment depends on the label of a segment far away. Recurrent neural networks such as LSTMs [15] have been used to model non-local dependencies where the model relies on the hidden state to remember the state of a token (e.g., a word) very far away. However, LSTMs are known to only remember a small number of tokens back, leading to rather “local” relationship learning [15, 30]. This drawback has lead to an increasing interest in the explicit modeling of non-local dependencies via pairwise interaction models such as transformers [30, 9, 8].

In a related line of work, graph convolutional networks (GCNs) have been proposed to model the pairwise dependencies of nodes in graph or 3D structured data such as citation networks and point clouds [24, 17, 32, 37]. This direct modeling of edges allows the network to learn non-local relationships. While typically viewed in its 1D sequential form, DNA can be represented as 3D genome structured data via Hi-C maps, as shown in Fig. 1(b). Hi-C maps are matrices that give the number of contacts between two segments of DNA, and normalized Hi-C maps tell us the likelihood of two locations interacting [2]. Using Hi-C data, segments of DNA can be represented as nodes on a graph, and edges are interactions between segements. Such interactions can be crucial in regulatory processes such as gene transcription [23]. That is, Hi-C contacts are a direct reflection of how distant epigenetic elements interact. We hypothesize that accounting for such interactions will lead to improved epigenetic state prediction accuracy.

In this work, propose ChromeGCN, a novel method that uses a fusion of both sequence and 3D genome data (in the form of Hi-C maps) to predict the epigenetic state of DNA segments. To the best of our knowledge, ChromeGCN is the first deep learning framework that successfully combines sequence and 3D genome data to model both local sequence features and long-range dependencies for epigenetic state prediction. ChromeGCN works by first representing DNA windows as a *d*-dimensional vector with a convolutional neural network on the local window sequence. We then revise the window vector using a graph convolutional network on all window relationships from Hi-C 3d genome data. We test ChromeGCN on datasets from two cell lines where we compare against the previous state of the art epigenetic state prediction methods. We demonstrate that ChromeGCN outperforms previous methods, especially for labels that are highly correlated with long-range chromatin interactions.

An important aspect of ChromeGCN is that it allows us to better understand Hi-C data in the context of epigenetic state. Hi-C maps tell us where the contacts are in the genome, but they don’t tell us important contacts for epigenetic labels. Using ChromeGCN, we propose Hi-C saliency maps to understand which Hi-C contacts are most important for epigenetic state labeling. Since ChromeGCN uses explicit long-range relationships from Hi-C data (as opposed to implicit long-range relationships using a recurrent neural network), we can easily understand the important relationships for greater interpretability.

The main contributions of this paper are:

1. We propose ChromeGCN, a novel framework that incorporates both local sequence and long-range 3D genome data for epigenetic state prediction.
2. We experimentally validate the importance of ChromeGCN on two cell lines from ENCODE, showing that modeling long range genome dependencies is critically important.
3. We introduce Hi-C saliency maps, a method to identify the important long range interactions for epigenetic state prediction from Hi-C data.

## 2 Background and Related Work

### 2.1 Predicting Epigenetic State Using Machine Learning

Computational models for accurately predicting epigenetic state labels from DNA sequence have gained popularity in recent years due to the urgency of the task for many applications. For instance, predicting how epigenetic effects vary when variants in DNA occur. The importance of computational modeling arises from the low cost and high speed in comparison to biological lab experiments.

One class of methods for state prediction used generative models in the form of position weight matrices (PWM) [29]. These methods construct motifs, or short contiguous sequences (often 8-20bp in length), which are representative of a particular epigenetic state label such as a transcription factor binding. A new sequence can then be classified according to how well it matches the motif. A significant drawback of using predefined motif features is that it is difficult to find the correct motifs for predicting unseen sequences [1]. Another class of methods use string kernels (SK) [10, 28], where some kernel function is built to capture the similarity between DNA segments according to substring patterns. However, these methods suffer from the issue of a predefined feature engineering. Moreover, these methods do not scale to a large number of sequences [36].

To overcome the issues of PWM and SK methods, researchers turned to automatic feature extraction using deep neural networks which have outperformed both generative PWM and SK methods [1, 36]. Convolutional neural networks (CNNs) were the first deep learning method to outperform previous methods. CNNs have been used extensively to learn features of DNA for sequence-based prediction [36, 1, 14, 19, 35]. The benefit of convolutional models is that they have an inductive bias for modeling translation invariant features in DNA sequences. This allows CNNs to effectively learn the correct “motifs” or kernels for chromatin labeling. There has since been several revisions to the original CNN models for marginally better feature extraction, such as adding a recurrent network on top of the CNN motif features [22, 20].

However, current state-of-the-art models only learn the features from the sequences of individual local windows and not between windows (i.e., longer-range interactions). Since DNA interacts with itself in the form of long-range 3D contacts, labeling the epigenetic states of a window can be affected by another distant window due. [16] use longer input sequences (131kbp), but the dependencies are modeled implicitly using dilated convolution across 128bp windows. Accordingly, methods that account for explicit long range 3D chromatin contacts are needed to model the true interactions in DNA.

### 2.2 DNA Interactions via Hi-C Maps

Hi-C experiments, and 3C experiments in general, are biological methods used to analyze the spatial organization of chromatin in a cell. These methods quantify the number of interactions between genomic loci. Two loci that are close in 3D space due to chromatin folding may be separated by up to millions of nucleotides in the sequential genome. Such interactions may result from biological functions, such as protein interactions [12]. Appendix Fig. 9 shows an example Hi-C map from the GM12878 cell line. The darker the lines indicate more DNA-DNA interactions.

Since the first Hi-C maps were generated, many works have been introduced to analyze the maps. [21] investigated the spatial relationships of co-localized TF binding sites within the 3D genome. They show that for certain TFs, there is a positive correlation of occupied binding sites with their spatial proximity in the 3D space. This is especially apparent for weak TF binding sites and at enhancer regions. [3] identified that the ZNF143 TF motif in the promoter regions provides sequence specificity for long range promoter-enhancer interactions. [34] identified coupling DNA motif pairs on long-range chromatin interactions. [25] use convolutional neural networks to predict Hi-C interactions from sequence inputs. None of the previous methods, however, use known Hi-C data to learn better feature representations of genomic sequences for epigenetic state prediction.

### 2.3 Graph Convolutional Networks

Graph convolutional networks (GCNs) were recently introduced to model non-local or non-smooth data [24, 7, 17, 13, 11, 31]. For the task of node classification, GCNs can learn useful node representations which encode both node-level features and relationships between connected nodes. Essentially, GCNs learn node representations by encoding local graph structures and node attributes, and the whole framework can be trained in an end-to-end fashion. Because of their effectiveness in learning graph representations, they achieve state-of-the-art results in node classification. The main assumption is that the input samples (in our case, individual DNA windows) are not independent. By modeling the graph dependency between samples, we can obtain a better representation of each of the samples. Non-local neural networks [32] are an instantiation of graph convolution, which was designed to model the long-range interactions in video frames.

## 3 Problem Formulation and Data Processing

The objective of epigenetic state prediction (i.e., chromatin effect prediction) is to tag segments of DNA with the probability of how likely a certain chromatin effect (aka epigenetic state label) is present. In our formulation, we define epigenetic state labels to include transcription factor (TF) binding, histone modifications (HM), and accessibility (DNase I). This is known as a multi-label classification task, where multiple labels can be positive at once (different from multi-class tasks where only one label can be positive). Formally, Given an input DNA window **x**_*i*_ (a segment of length *T*), we want to predict 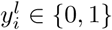 for a label *l*, where *l* ranges from 1 to *L*.

### Sequence Data

We derive epigenetic labels using ChIP-seq data from ENCODE [6]. We use the cell lines GM12878 and K562, two of the most widely used from ENCODE and Roadmap [6, 18]. For each cell line, we use all windows which have at least one positive epigenetic ChIP-seq peak. We consider any peak from ENCODE to be a positive peak. We follow a similar setup as in [36] where we bin the DNA into 1000bp windows. If any ChIP-seq peak overlaps with at least 100bp of a particular window, we consider that a positive window for that chromatin label. We then extract the 2000bp sequence surrounding the center of each window as the input features, as done in [35], since the motif for a particular signal may not be contained fully in the 1000bp length window. Although we use the 2000bp sequence, we consider each window to be the original non-overlapping 1000bp for notation purposes. An illustration of how sequences are extracted is shown in Appendix Fig. 10 (a). Following [36], we use chromosome 8 for testing and also add chromosomes 1 and 21. Chromosomes 3, 12, and 17 are used for validation, and all other chromosomes (excluding X and Y) are used for training. The datasets are summarized in Table 1.

**Table 1:** Datasets Summary. GM12878 contains 103 total epigenetic state labels, and K562 contains 164 total labels. We use the same chromosomes for training, validation, and testing for both datasets.

| Cell Line | Train Windows | Valid Windows | Test Windows | TFs | HMs | DNase I |
| --- | --- | --- | --- | --- | --- | --- |
| GM12878 | 368,082 | 89,911 | 79,731 | 90 | 11 | 2 |
| K562 | 457,609 | 106,777 | 117,815 | 150 | 12 | 2 |

### 3D Genome Data

We then use 3D genome data from Hi-C contact maps to extract interaction evidence between the DNA windows. We use 1000bp resolution intra-chromosome Hi-C data from [23] (for K562, the lowest resolution is 5000bp, so we upsample to get 1000bp resolution). We convert the Hi-C contact map for each chromosome into a graph whose nodes are 1000bp DNA windows and whose edges represent contact between two 1000bp windows. Since the full Hi-C contact for each chromosome is too dense, we rank each contact edge, and use the top 500,000 Hi-C contacts as edges per chromosome (each chromosome maps to a Hi-C graph). Contacts are ranked using the *SQRTVC* normalization from [23], which normalizes for the distance between two positions so that long-range contacts are included in the top 500k.

## 4 Method

Our goal is to learn a model *f* which takes in a DNA subsequence window **x**_*i*_ and predicts the probability of a set of epigenetic labels **ŷ**_*i*_ = *f* (**x**_*i*_), where **ŷ**_*i*_ is an *L* dimensional vector. Our method, ChromeGCN uses three submodules for *f*: *f*_*CNN*_, *f*_*GCN*_, and *f*_*Pred*_. The first module, *f*_*CNN*_, models local sequence patterns from each window using a convolutional neural network. This module takes as input **x**_*i*_ and outputs a vector representation **h**_*i*_ = *f*_*CNN*_ (**x**_*i*_). The second module, *f*_*GCN*_, models long range 3D genome dependencies between windows using a graph convolutional network. This module takes as input all window vectors **h**_*i*_ concatenated as **H**, as well as their Hi-C relationships represented by adjacency matrix **A**, and outputs refined representations of all windows **Z** = *f*_*GCN*_ (**H, A**). The **z**_*i*_ of each window now encodes both window sequence patterns and the relationships between windows. We can then predict the epigenetic labels using a classifier function on each **z**_*i*_ using **ŷ**_*i*_ = *f*_*Pred*_(**z**_*i*_). An overview of ChromeGCN is shown in Fig. 1(c). The following subsections explain each submodule in detail.

### 4.1 Modeling Local Sequence Representations Using Convolutional Neural Networks

Following the recent successes in many epigenetic label prediction tasks [36, 1, 22, 20, 35], we learn a representation of each genomic window sequence **x**_*i*_ using a convolutional neural network (CNN). CNNs have become the de facto standard for encoding short DNA windows due to their properties, which effectively capture local sequence structure. Each learned kernel, or filter, in CNNs effectively learns a DNA “motif”, or short contiguous sequence representative of a particular output label [1]. Since many epigenetic processes are hypothesized to be dependent on motifs [3], CNNs are a good choice for encoding DNA.

This module, *f*_*CNN*_, takes an input genomic sequence window **x**_*i*_, and outputs an embedding representation vector **h**_*i*_. We represent window **x**_*i*_ of length *τ* as a one-hot encoded matrix 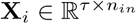, where *n*_*in*_ is 4, representing the base-pair characters A,C,G, and T. Convolution with filters (i.e. learned motifs) of length *k < τ* takes an input data matrix **X**_*i*_ of size *τ* × *n*_*in*_, and outputs a matrix **X**′_*i*_ of size *τ* × *n*_*out*_, where *n*_*out*_ is the chosen dimension of the learned hidden representations:

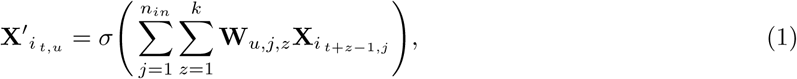

where 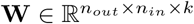 are the trainable weights, and *σ* is a function enforcing element-wise nonlinearity. Eq. 1 can then be repeated for several layers where each successive layer uses a new **W** and *n*_*in*_ is replaced with *n*_*out*_ from the previous layer. In our implementation, we use six layers of convolution where each successive layer learns higher-order motifs of the window. After the convolutional layers, the output of the last layer is flattened into a vector and then linearly transformed into a lower-dimensional vector of size *d*, which we denote **h**_*i*_. Succinctly, the CNN module computes the following: **h**_*i*_ = *f*_*CNN*_ (**x**_*i*_) for each window.

### 4.2 Modeling Long-Range 3D Genome Relationships Using Graph Convolutional Networks

While CNN models work well on independent local window sequences, they disregard known long-range relationships between windows that are influential in the epigenetic state. One option would be to extend the window size. However, due to the 3D shape of DNA, long-range contact dependencies may be located millions of base-pairs apart, making current convolutional models infeasible. In this subsection, we introduce the *f*_*GCN*_ module, a method to explicitly and efficiently model such long-range interactions using graph convolutional networks.

Known long-range relationships in the 3D genome are available in the form of Hi-C contact maps. A Hi-C map can be represented as an adjacency matrix **A**, where the nonzero elements indicate contacts in the 3D space between two DNA windows^2^. In our formulation, we represent sequence windows **x**_*i*_ as nodes on a graph, and **A** are the edges between the windows. We can then use a modified version of graph convolutional networks [17] (GCN) to update each **x**_*i*_ with its neighboring windows **x**_*j*_, *j* ∈ 𝒩 (*i*), where 𝒩 (*i*) denotes the neighbors of node *i* obtained from **A**. The GCN works by revising a window’s representation 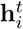, where 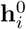 is from the output of the first module, *f*_*CNN*_. Specifically, each 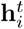 is revised using a parameterized summation of neighbors, 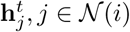:

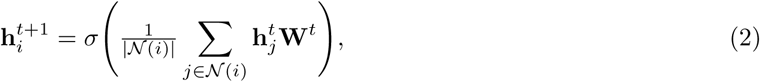

where *σ*(·) is a non-linear activation function such as tanh and **W**^*t*^ ∈ ℝ^*d*×*d*^ is a linear feature transformation matrix for the GCN layer *t*. Importantly, using the summation in Eq. 2, the representation of each DNA window **x**_*i*_ is updated based on the representation of its neighbors (windows that interact with **x**_*i*_ in the 3D genome). We can compute the simultaneous update of all windows together by concatenating all **h**_*i*_ denoted **H**^*t*^ ∈ ℝ^*N* ×*d*^ where *N* is the number of DNA windows and *d* is the dimension of each **h**_*i*_. The simultaneous update can then be written as:

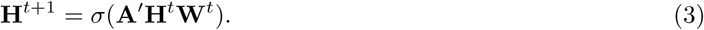

where 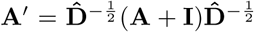, is the normalized adjacency matrix and 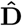 is the diagonal degree matrix of (**A** + **I**).

In our experiments, we use a variant of the graph convolutional network, which uses a gating function allowing the model to use or not use neighboring windows to update each **h**_*i*_:

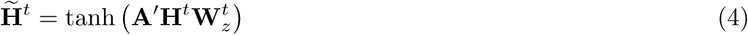

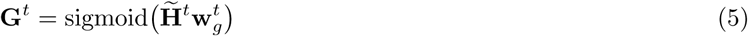

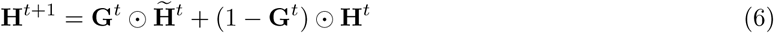

where 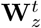 is a linear transformation matrix, and 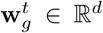 is used to compute the gate. **G**^*t*^ allows the model to selectively choose between using the neighborhood representation of nodes, 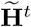, or the independent representation, **H**^*t*^. ⊙ is an Hadamard product. Equations 3-6 indicate one layer update of GCN window embeddings **H**. In our experiments, we use a 2-layer gated GCN, and we denote the final output of all **H**^*t*^ as **Z**, where vector **z**_*i*_ of **Z** represents the output of window *i*. In summary, the GCN module computes the following: **Z** = *f*_*GCN*_ (**H, A**).

### 4.3 Predicting Label Probabilities for Each Window

After **Z** is computed, we then use a linear classifier layer, *f*_*Pred*_ to classify each **z**_*i*_ into its output space (a set of epigenetic labels): 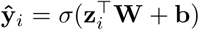. In summary, the prediction **ŷ**_*i*_ for a particular input sample **x**_*i*_ can be decomposed as three steps:

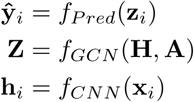

### 4.4 Identifying and Interpreting Important Hi-C Edges via Hi-C Saliency Maps

One benefit of the ChromeGCN formulation is that we use explicit long-range window dependencies for epigenetic state prediction. As a result, we propose a method to identify the important dependencies for ChromeGCN’s predictions. We call our proposed method Hi-C saliency maps. Saliency maps were introduced by [27] to understand the importance of each pixel in an input **x**_*i*_ for the prediction of the image’s true class. We instead are trying to understand the importance of each edge in **A**′ for the prediction of the epigenetic state over all windows. The **A**′ saliency map is defined as the absolute value gradient of a true class prediction 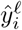 with respect to **A**′, where *ℓ* is a true class for sample *i*. The absolute value gradient is then element-wise multiplied by **A**′ to zero out “non-edges”. Since there are *N* samples, and there can be multiple true labels *ℓ* for a particular sample **ŷ**_**i**_, we define the Hi-C saliency map, *S*_*Hi*-*C*_, as the accumulated absolute gradient over all samples and true labels w.r.t **A**′:

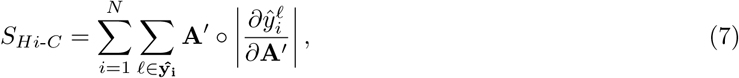

where ○ is the Hadamard product. Since the saliency map of all windows *N* are accumulated, we normalize *S*_*Hi*-*C*_ across each row to a 0-1 range so that we can interpret the edges at each window.

While Hi-C contact maps tell us *where* the contacts are, Hi-C saliency maps show us *how important* each contact is for the epigenetic states. We define Eq. 7 to be over all labels, but we can easily visualize the Hi-C saliency, or important edges for one particular label. We show both the full Hi-C saliency across all labels, as well as for one specific label (YY1) in the experiments.

### 4.5 Model Variations

To test the effectiveness of the long range dependencies, we use the following ChromeGCN variations. Each variation uses the same model, with different edge dependencies in the form of **A**.

#### ChromeGCN_*const*_

Instead of using Hi-C edges we use a constant set of nearby neighbors according to the 1D sequential DNA representation. We define each window **x**_*i*_’s neighbors to be the windows surrounding **x**_*i*_ (7 on each side: **x**_*i*−7_, …, **x**_*i*+7_) which we denote as **A**_*const*_. **Z** = *GCN* (**A**_*const*_, **H**). This variant allows us to see whether the very long range interactions from the normalized Hi-C maps are useful.

#### ChromeGCN_*Hi-C*_

This variation uses the original Hi-C adjacency matrix, **A**_*Hi*-*C*_. **Z** = *GCN* (**A**_*Hi*-*C*_, **H**). Since we used use the top 500k Hi-C edges using a the SQRTVC normalization, many of these edges are far away in the 1D space.

#### ChromeGCN_*const*+*Hi-C*_

Lastly, we use a combination of the constant neighborhood around each window and the Hi-C adjacency matrix, which integrates close and far windows for each window. This varition uses the following function: **Z** = *GCN* (**A**_*const*+*Hi*-*C*_, **H**).

### 4.6 Model Details and Training

To circumvent GPU memory constraints of training end-to-end, we pretrain the *f*_*CNN*_ model by classifying each **h**_*i*_ with the classification function **ŷ**_*i*_ = *f*_*Pred*_(**h**_*i*_). Once the pretraining converges on the training set, we use the trained weights **h**_*i*_ for each sample as fixed inputs to *f*_*GCN*_. While we pretrain *f*_*CNN*_, ChromeGCN is still end-to-end differentiable, making it possible to use sequence visualization methods such as DeepLIFT [26] for a particular window.

For all model predictions, we run the forward and the reverse complement through simultaneously and average the output of the two. All DNA window inputs are encoded using a lookup table that maps each character A, C, G, T, and N (unknown) to a *d*-dimensional vector. The output of the encoding is a *d* × *τ* matrix, where*τ* denotes sequence length (*τ* = 2000 in our experiments).

All of our models are trained using stochastic gradient descent with momentum of 0.9 and a learning rate of 0.25. The CNN model is trained using a batch size of 64, and the GCN and RNN models are trained using an entire chromosome as a batch (since each is modeling the between window dependencies of an entire chromosome at once). The CNN model projects each window to a vector of dimension 128. The GCN uses two layers of feature dimension 128 at each layer.

ChromeGCN predicts the probabilities of all labels for each window: **ŷ**_*i*_ ∈ ℝ^*L*^, where *L* is the total number of labels. For our loss function, we use the mean binary cross-entropy across all samples *N* and labels *L*:

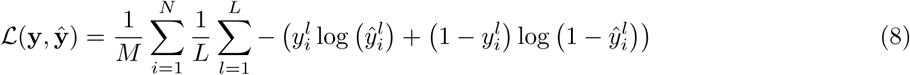

## 5 Experiments and Results

### 5.1 Baselines

We compare against the state-of-the-art epigenetic state prediction model from [35], referred to as the CNN baseline, as well as the recurrent model from [22]. Since our model outputs labels for TFBS, HMs, and accessibility, motif-based methods [4] aren’t applicable. Since we have 368,082 training samples, kernel-based methods such as [10] aren’t applicable. [36] compared their CNN to a modified version of [10], which only used a small number of training samples, and the CNN model was significantly better. Our CNN baseline, [35] is an improved version from [36]. Furthermore, the focus of our study is to show that state-of-the-art deep learning models are missing important long range dependencies in the genome.

#### CNN [35]

To illustrate the importance of the GCN, we compare against the outputs from the *f*_*CNN*_ module: **ŷ**_*i*_ = *f*_*Pred*_(**h**_*i*_). This is the 6-layer CNN model from [35] (we modify the last layer in order to extract a *d* dimensional feature vector output). This is the same CNN that is pretrained for ChromeGCN to produce each **h**_*i*_.

#### DanQ [22]

This model uses a recurrent neural network (RNN) on top of CNN outputs within a window. It still uses local sequence window inputs, but models relationships between sequence patterns via an LSTM.

#### ChromeRNN

As a baseline to compare against using GCNs for long range dependency modelling, we construct an RNN model on the window embeddings **h**_1_, **h**_2_, …, **h**_*N*_. After pretraining the CNN module *f*_*CNN*_, The RNN model takes in all window embeddings at once and models the sequential dependencies among windows: (**z**_1_, **z**_2_, **z**_3_, …, **z**_*N*_) = *f*_*RNN*_ (**h**_1_, **h**_2_, **h**_3_, …, **h**_*N*_). As with ChromeGCN, the RNN is shared across chromosomes, but does not cross chromosomes. In other words, the embeddings are updated one chromsome at a time *f*_*RNN*_ (**h**_*i*_, **h**_*i*+1_, …, **h**_*C*_) where *C* is the total number of windows for a chromosome. We note this is different from the DanQ baseline [22] which uses an RNN *within* windows [22]. ChromeRNN instead is for modeling dependencies *between* windows. In our experiments, we use an LSTM [15] with the same number of layers and hidden units as the GCN.

### 5.2 Prediction Performance

To evaluate the methods, we use area under the ROC curve (AUROC), area under the precision-recall curve (AUPR), and the Mean Recall at 50% False Discovery Rate (FDR) cutoff. Table 2 shows the mean metric results across all epigenetic labels for each cell line. Modeling long-range dependencies results in significant improvements over the baseline CNN model, which does not account for such long range interactions. For instance, with respect to AUPR for GM12878, ChromeRNN improves upon the CNN from 0.350 to 0.384, and ChromeGCN outperforms ChromeRNN to achieve a mean AUPR of 0.395. Also, we see that the ChromeRNN outperforms DanQ, indicating that using an RNN to model between window dependencies is more important than within window features. Moreover, we can see that the ChromeGCN_*Hi*-*C*_ models out-perform ChromeRNN, indicating that not only the closely neighboring windows in the 1D space contribute to the improvements, but also the close neighbors in the 3D space, as indicated by the used Hi-C maps. ChromeGCN outperforms the baselines on the TF and DNase I labels, and ChromeRNN outperforms all other methods on the HM labels. This indicates that non-local modeling is particularly important for TF binding and accessibility. We provide the performance results of each label type (TF, HM, DNase I) are shown in Appendix Tables 3 and 4. We also provide detailed plots of ROC and Precision-Recall curves in Appendix Figures 5, 6, 7, and 8.

**Table 2:** Performance results. For both cell lines, GM12878 and K562, we show the average across all labels for three different metrics. Our method, using a graph convolutional network (GCN) to model long range dependencies helps improve performance over the baseline CNN model which assumes all DNA segments are independent. Detailed results of all cell lines and label types are shown in the Appendix.

|  | GM12878 |  |  | K562 |  |  |
| --- | --- | --- | --- | --- | --- | --- |
|  | Mean AUROC | Mean AUPR | Mean Recall at 50% FDR | Mean AUROC | Mean AUPR | Mean Recall at 50% FDR |
| CNN [35] | 0.895 | 0.350 | 0.293 | 0.894 | 0.325 | 0.265 |
| DanQ [22] | 0.886 | 0.348 | 0.290 | 0.900 | 0.343 | 0.290 |
| ChromeRNN | 0.906 | 0.384 | 0.342 | 0.910 | 0.365 | 0.327 |
| ChromeGCN<sub>const</sub> | 0.904 | 0.377 | 0.331 | 0.904 | 0.358 | 0.321 |
| ChromeGCN<sub>Hi-C</sub> | 0.904 | 0.385 | 0.341 | 0.903 | 0.358 | 0.319 |
| ChromeGCN<sub>const+Hi-C</sub> | **0.909** | **0.395** | **0.356** | **0.912** | **0.372** | **0.338** |

Furthermore, since ChromeGCN models the window relationships explicitly and not recurrently, we obtain a significant speedup at test time over the ChromeRNN baseline. ChromeGCN achieves a 6.3x speedup on the three GM12878 test chromosomes, and 6.8x speedup at test time on the three K562 test chromosomes.

### 5.3 Analysis of Using Hi-C Data

Fig. 2 shows a a detailed comparison of ChromeGCN_*Hi*-*C*_ vs the baseline CNN model across three different metrics for both GM12878 and K562. Each point represents a label, and the y-axis shows the absolute improvement of the ChromeGCN_*Hi*-*C*_ model over the CNN. The labels are sorted on the x-axis by the average degree of the label’s positive samples (i.e., windows where the label is positive) on the Hi-C map. We can see that for all three metrics, the improvements of the ChromeGCN over the CNN increase as the average degree of the labels increase. This indicates that the ChromeGCN is important for labels that have many neighbors in the Hi-C graph (i.e., those that are frequently in contact with other segments in the 3D space). Two of the transcription factors which obtain the highest performance increase (in the top 5) from using ChromeGCN over CNN, CEBPB STAT3, are validated by [21], which show that these two TFs commonly co-occur with other TFs in the 3D space when binding.

**Fig. 2:**
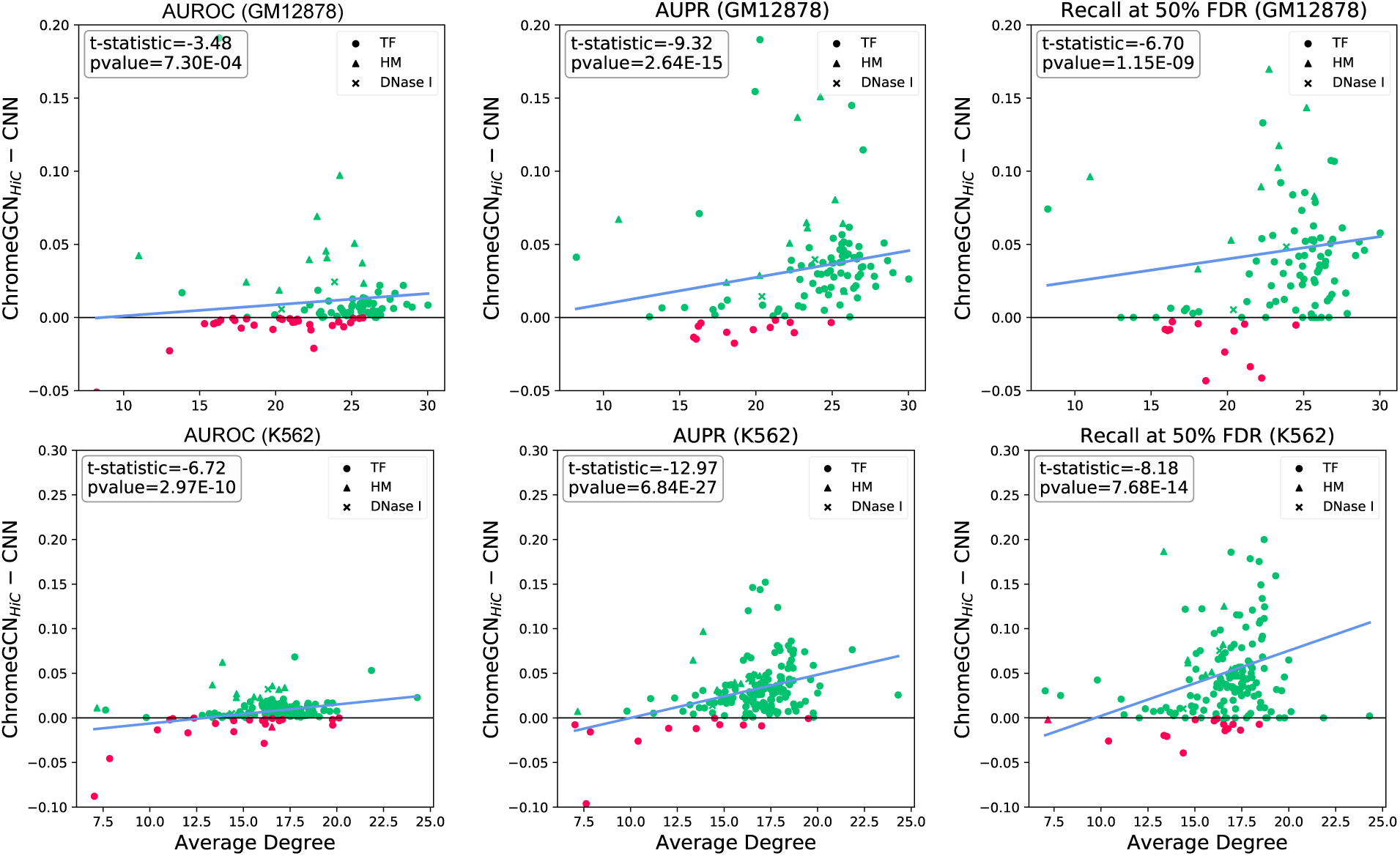
Comparison of our ChromeGCN_*Hi*-*C*_ method vs the baseline CNN [35] for 3 Metrics. Each point represents one epigenetic state label. The labels are sorted in the x-axis by the average degree of their positive windows. The y-axis indicates absolute increase of the ChromeGCN_*Hi*-*C*_ over the CNN model. As the average degree increases, the improvement of the ChromeGCN_*Hi*-*C*_ model increases over the CNN. Green points indicate ChromeGCN_*Hi*-*C*_ performed better, red indicate the CNN performed better. The blue line shows the linear trend line. The ChromeGCN_*Hi*-*C*_ is significantly better, as demsonstrated by the pvalues from a pairwise t-test.

The p-values shown are computed by a pairwise t-test across all labels. The ChromeGCN_*Hi*-*C*_ model significantly outperforms the CNN model in all three metrics. Importantly, our results indicate that by using the long-range interactions given by Hi-C data, we can obtain improvements in modeling the chromatin state labeling, resulting in better classification accuracy.

### 5.4 Long Range Interaction Visualization

A significant merit of ChromeGCN is that by using known 3D genome relationships, we can find and visualize the critical relationships for epigenetic state prediction. To understand how important the Hi-C edges are for the predictions of ChromeGCN, we visualize the saliency map of **A**′, as explained in Section 4.4.

Fig. 3 shows the Hi-C saliency map for chromosome 8 in GM12878. Fig. 3 (left) shows all 500k Hi-C contacts used chromosome 8. Windows are represented as points along the circle, with a total of 23,600 windows. Lines between the windows represent Hi-C edges, and the darkness of the line represents the saliency, or importance of that edge for chromatin state prediction across all windows in chromosome 8. Fig. 3 (right) shows the saliency map for 250 windows (total of 250kbp input) in chromosome 3. Cell (*i, j*) tells us the importance of window column *j* for the prediction of window row *i* labels.

**Fig. 3:**
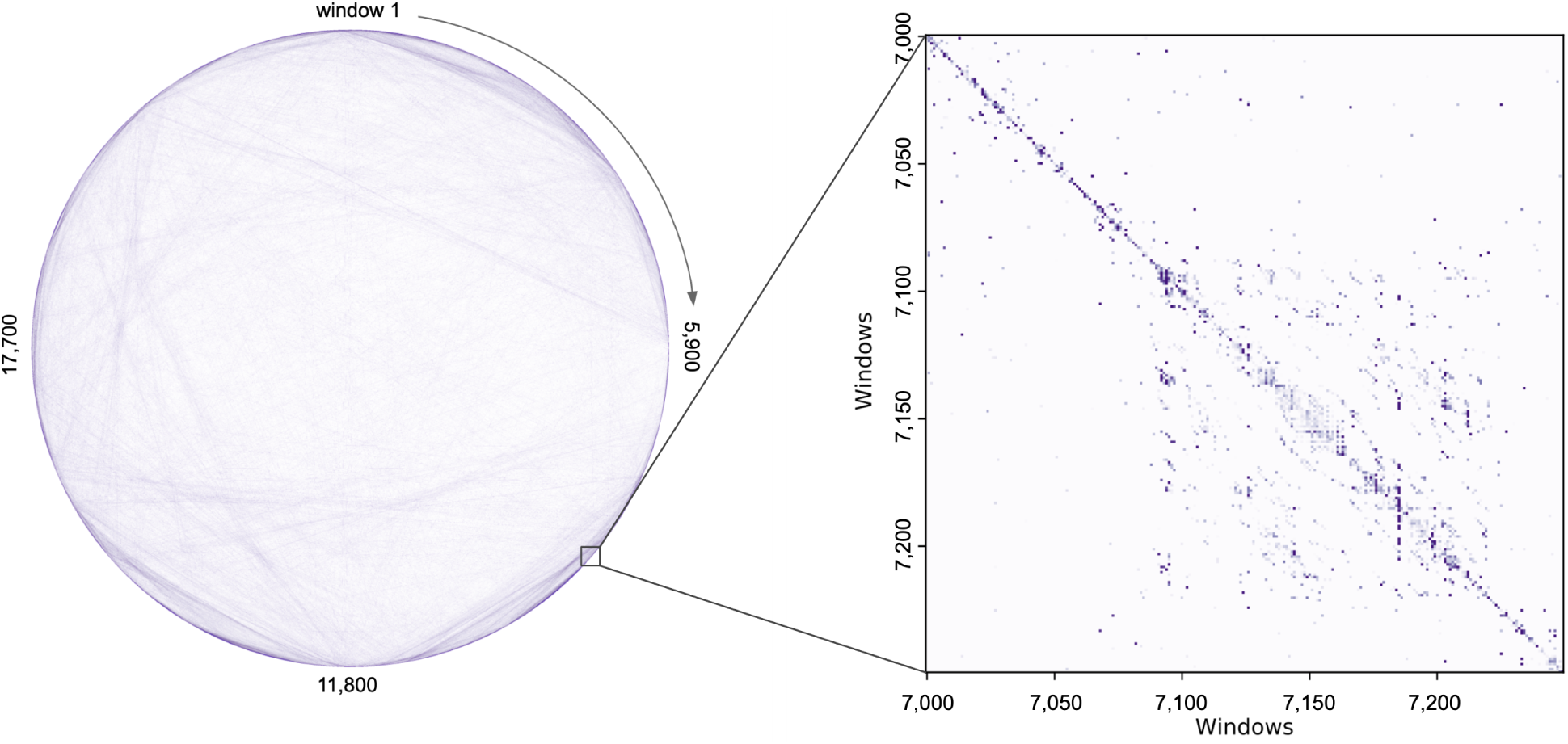
Hi-C Saliency Map Visualization. *Left:* Saliency Map for the all 500k edges in **A**_*Hi*-*C*_ for GM12878 Chromosome 8 (total of 23,600 windows). The darker the line, the more important that edge was for predicting the correct Epigenetic state, indicating that the Hi-C data was used by the GNN for that particular interaction. *Right:* Fine grained analysis of the Chromosome 8 Saliency Map. This figure shows the normalized Saliency Map values for for 250 windows (total of 250kbp input) in chromosome 8.

While Fig 3 shows the Hi-C saliency map for all epigenetic state labels, we can also visualize the Hi-C saliency map for individual labels. The inner loop of Eq. 7 changes to only use the label of interest. In Appendix Fig 4, we show the Hi-C saliency map for the YY1 transcription factor on GM12878 chromosome 8. This gives us insight into the important 3D contacts for YY1 binding.

## 6 Conclusion

In this work, we present ChromeGCN, a novel framework that combines both local sequence and long-range 3D genome data (via Hi-C data) for epigenetic state prediction. We show experimentally that ChromeGCN outperforms previous state-of-the-art methods that only use local sequence data. Additionally, we show that we can identify and visualize the important 3D genome dependencies using the proposed Hi-C saliency maps. We plan to further investigate the value of Hi-C saliency maps in future work.

While we demonstrate the importance of ChromeGCN on the task of epigenetic state prediction, ChromeGCN is a generic model for incorporating 3D genome structure into any genome sequence prediction task. We hope that this work encourages researchers to use known long-range relationships from 3D genome data when constructing machine learning models. We plan to release our data and code for greater visibility. ChromeGCN introduces an effective and efficient framework to model such relationships for better chromatin modeling, as well as an easy way to interpret important relationships.

## 7 Appendix

### 7.1 YY1 Hi-C Saliency Map

**Fig. 4:**
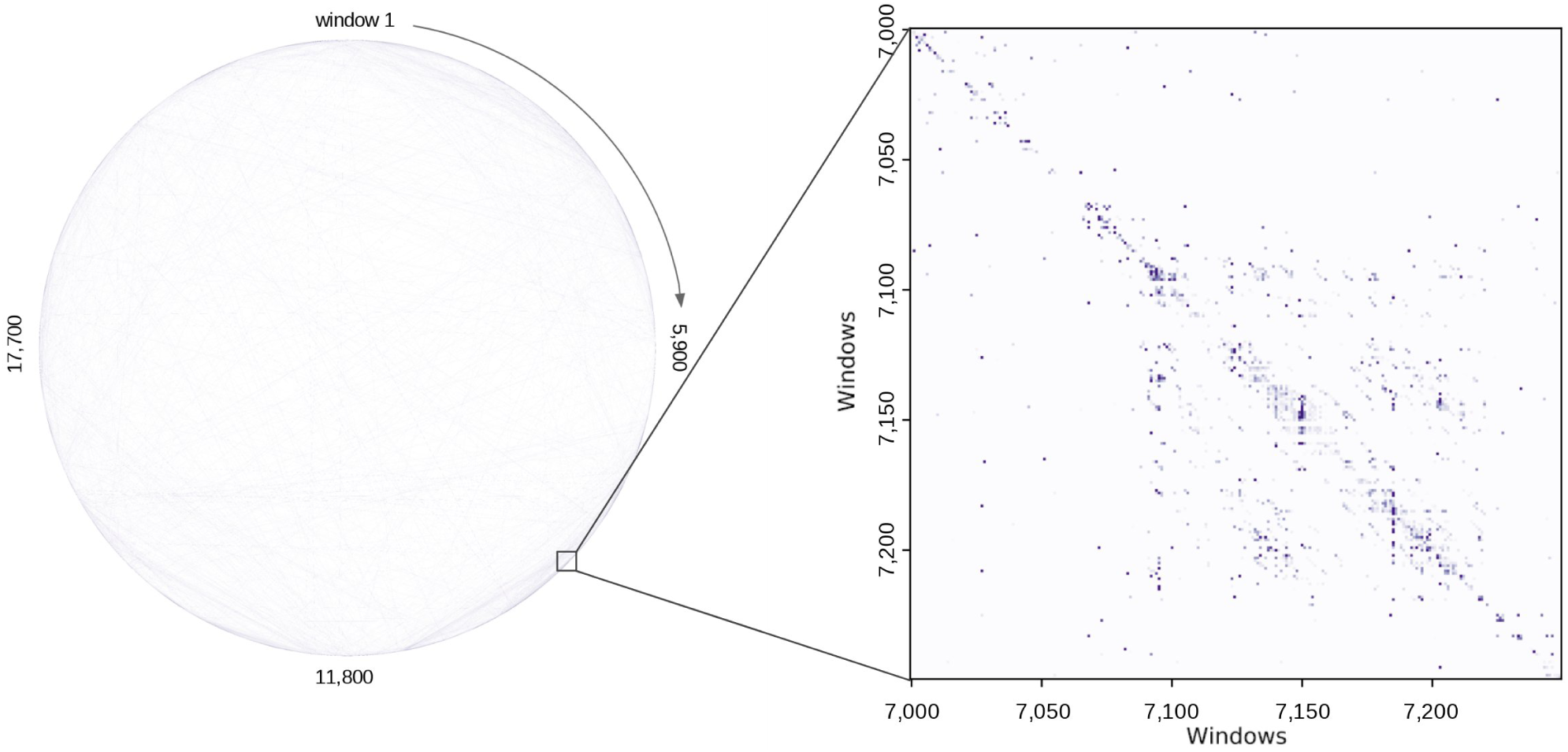
Hi-C Saliency Map for the YY1 transcription factor on GM12878 Chromosome 8.

### 7.2 Additional Performance Tables and Figures

**Table 3:**
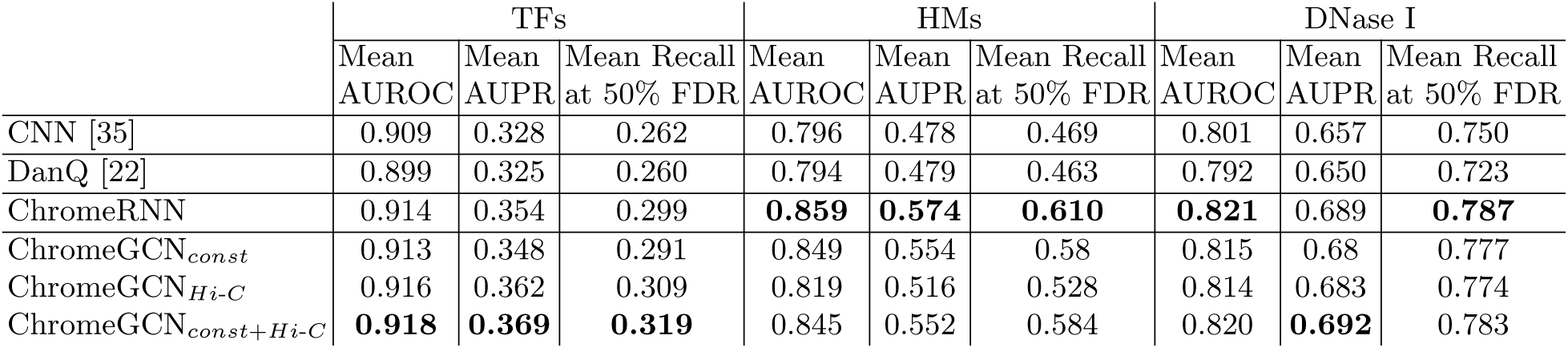
Performance on GM12878 for each label category.

**Table 4:** Performance on K562 for each label category.

|  | TFs |  |  | HMs |  |  | DNase I |  |  |
| --- | --- | --- | --- | --- | --- | --- | --- | --- | --- |
|  | Mean AUROC | Mean AUPR | Mean Recall at 50% FDR | Mean AUROC | Mean AUPR | Mean Recall at 50% FDR | Mean AUROC | Mean AUPR | Mean Recall at 50% FDR |
| CNN [35] | 0.905 | 0.314 | 0.256 | 0.78 | 0.410 | 0.317 | 0.792 | 0.625 | 0.683 |
| DanQ [22] | 0.909 | 0.331 | 0.279 | 0.796 | 0.438 | 0.365 | 0.797 | 0.638 | 0.693 |
| ChromeRNN | 0.917 | 0.349 | 0.305 | **0.848** | **0.520** | **0.528** | 0.817 | 0.664 | 0.742 |
| ChromeGCN<sub>const</sub> | 0.911 | 0.342 | 0.301 | 0.84 | 0.507 | 0.507 | 0.810 | 0.657 | 0.729 |
| ChromeGCN<sub>Hi-C</sub> | 0.912 | 0.346 | 0.306 | 0.807 | 0.454 | 0.412 | 0.812 | 0.657 | 0.730 |
| ChromeGCN<sub>const+Hi-C</sub> | **0.919** | **0.358** | **0.320** | 0.837 | 0.495 | 0.484 | **0.821** | **0.67** | **0.752** |

**Fig. 5:**
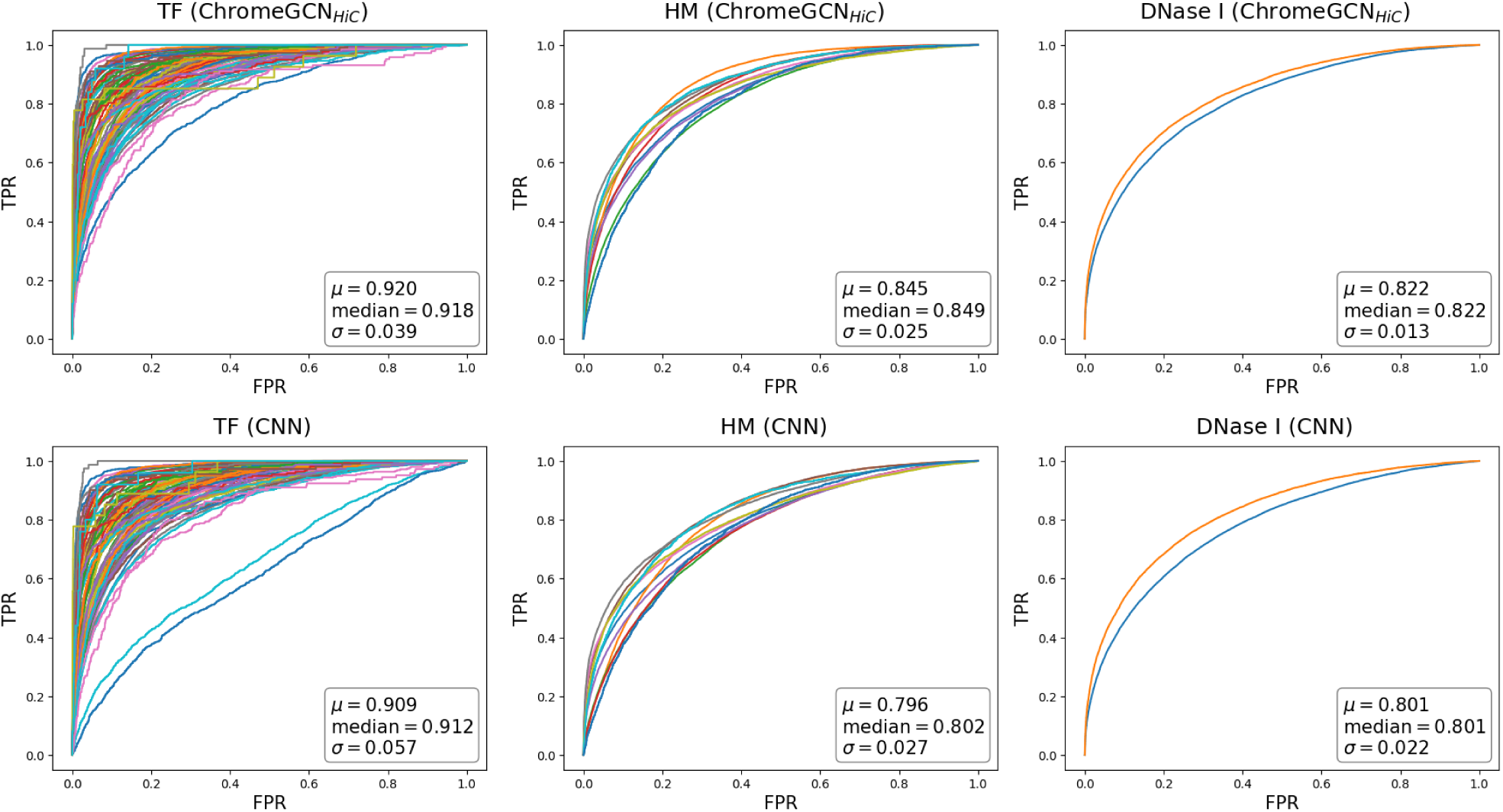
ROC curves for GM12878. The top row shows the ChromeGCN_*Hi*-*C*_ variant, and the bottom row shows the CNN[35]. The columns are divided into the 3 types of labels: transcription factors (TFs), histone modificiations (HMs), and DNA accesibility (DNase I). The color of each curve represents a different label, where they are consistent across columns. The box in each plot shows the statistics of the area under the curves (AUC). ChromeGCN outperforms the CNN for all Epigenetic state labels.

**Fig. 6:**
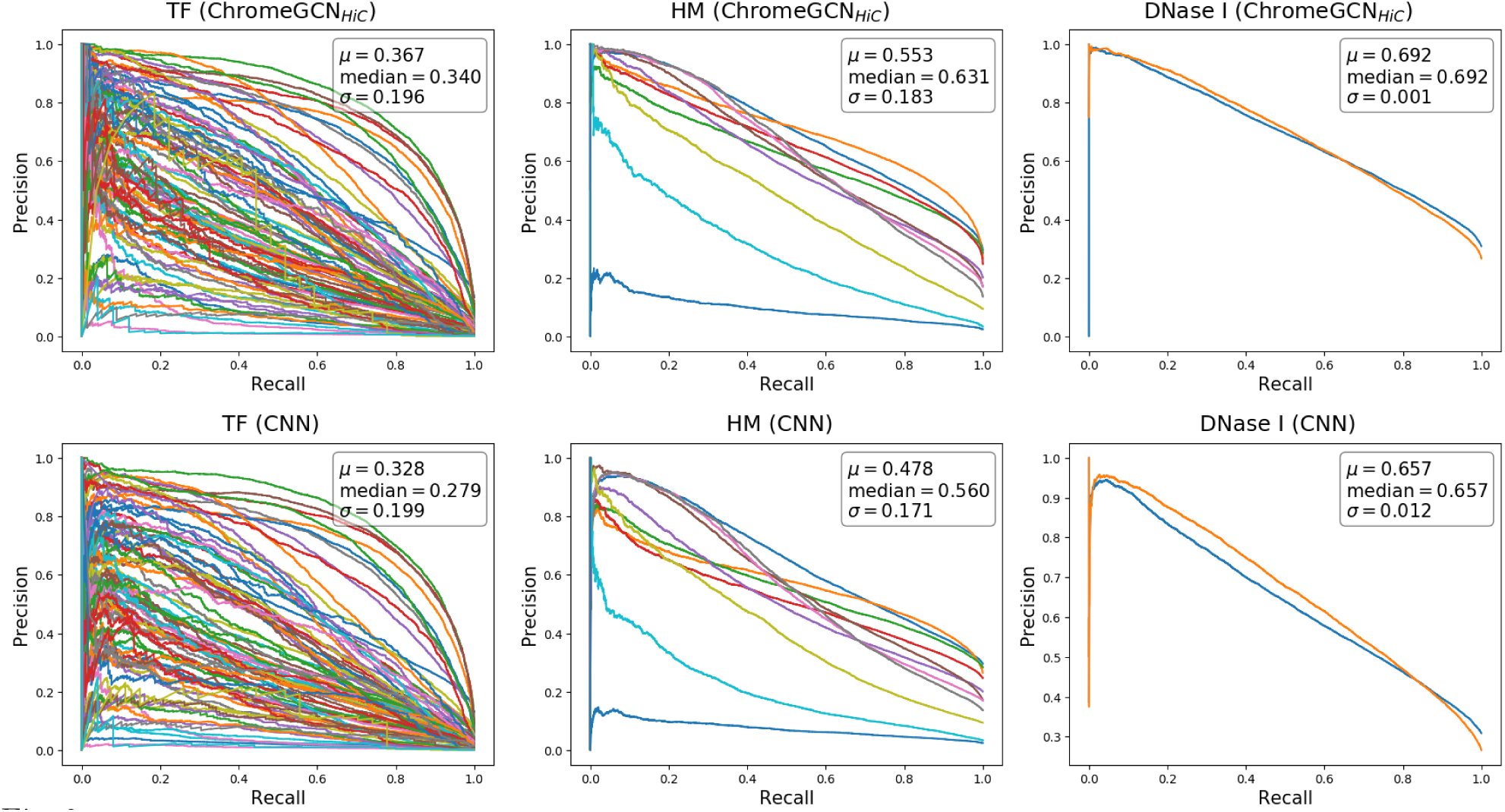
Precision-Recall curves for GM12878. The top row shows the ChromeGCN_*Hi*-*C*_ variant, and the bottom row shows the CNN[35]. The columns are divided into the 3 types of labels: transcription factors (TFs), histone modificiations (HMs), and DNA accesibility (DNase I). The color of each curve represents a different label, where they are consistent across columns. The box in each plot shows the statistics of the area under the curves (AUC). ChromeGCN outperforms the CNN for all Epigenetic state labels.

**Fig. 7:**
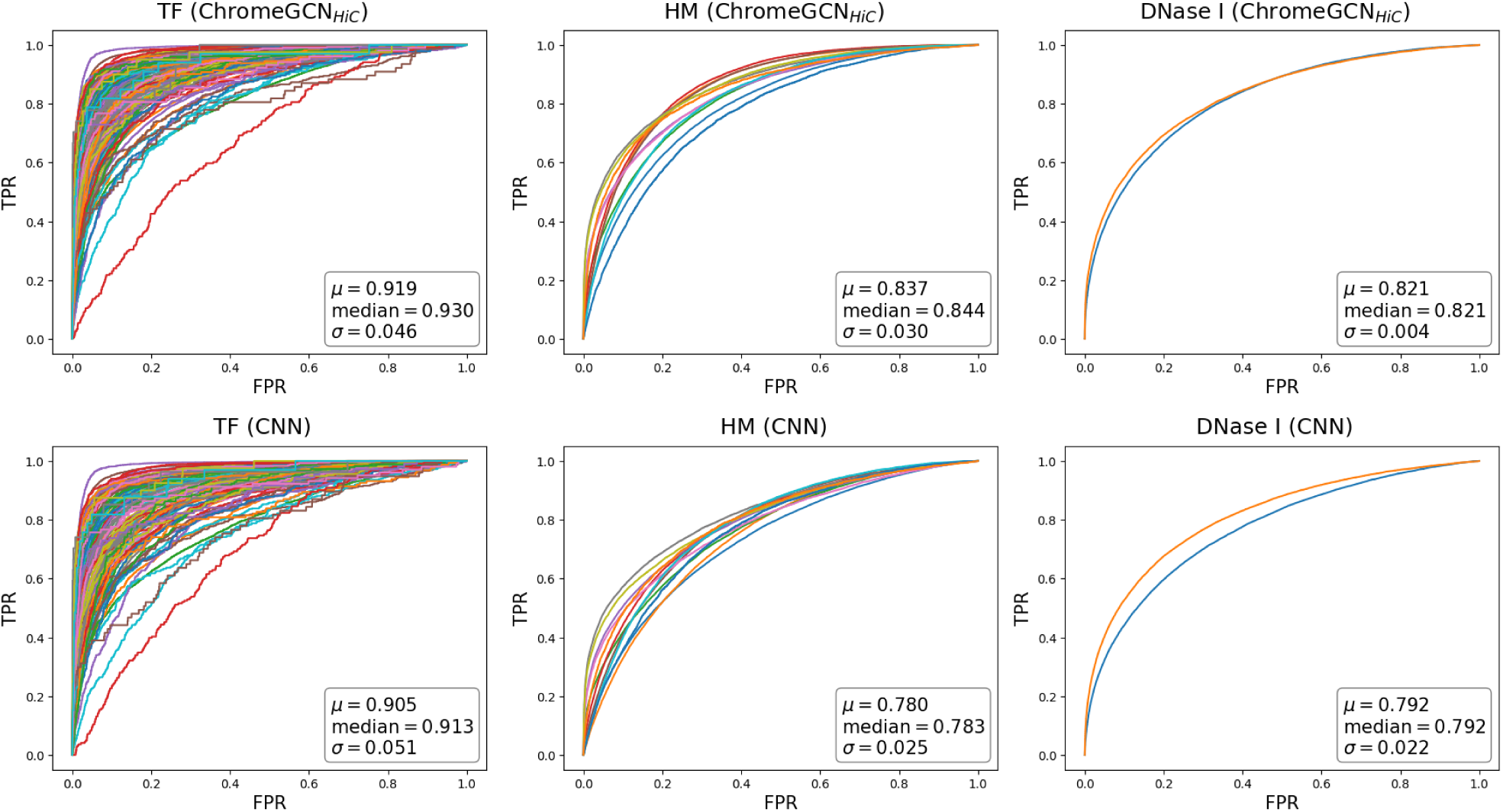
ROC curves for K562. The top row shows the ChromeGCN_*Hi*-*C*_ variant, and the bottom row shows the CNN[35]. The columns are divided into the 3 types of labels: transcription factors (TFs), histone modificiations (HMs), and DNA accesibility (DNase I). The color of each curve represents a different label, where they are consistent across columns. The box in each plot shows the statistics of the area under the curves (AUC). ChromeGCN outperforms the CNN for all Epigenetic state labels.

**Fig. 8:**
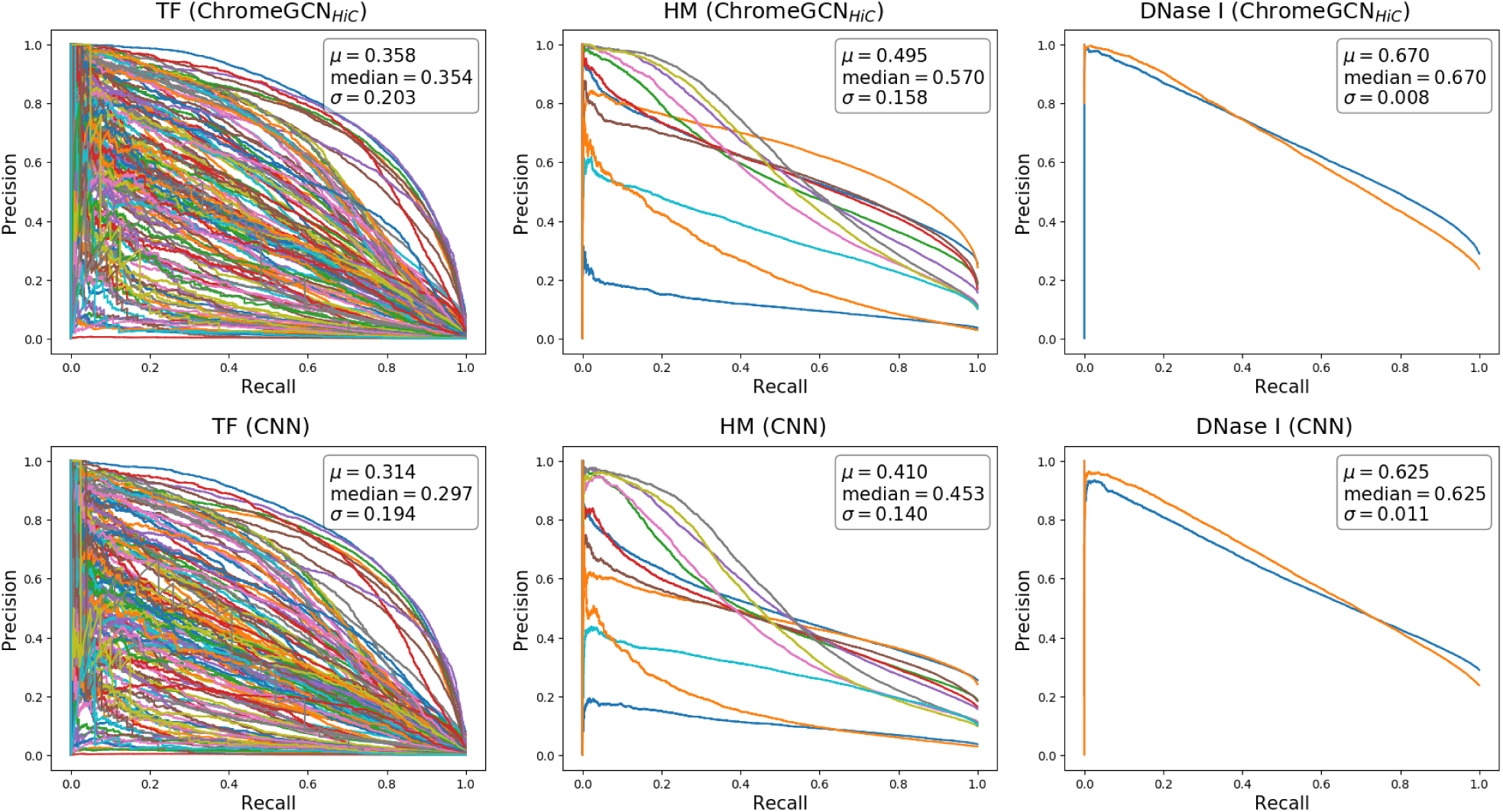
Precision-Recall curves for K562. The top row shows the ChromeGCN_*Hi*-*C*_ variant, and the bottom row shows the CNN[35]. The columns are divided into the 3 types of labels: transcription factors (TFs), histone modificiations (HMs), and DNA accesibility (DNase I). The color of each curve represents a different label, where they are consistent across columns. The box in each plot shows the statistics of the area under the curves (AUC). ChromeGCN outperforms the CNN for all Epigenetic state labels.

### 7.3 Data Details

**Fig. 9:**
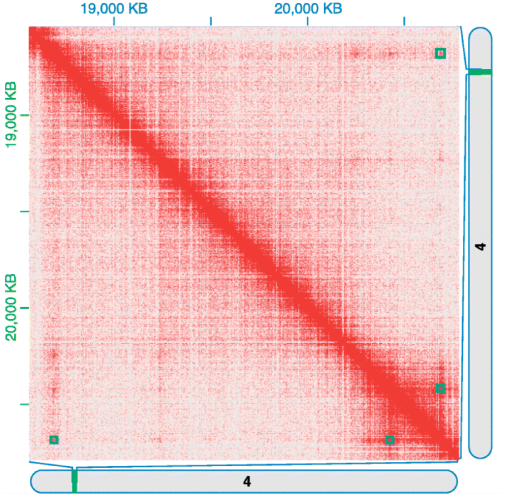
Example Hi-C map showing the (chr4:18000000-21000000) region from GM12878 cell line [33]. Red blocks indicate DNA interactions, the darker red indicates more interactions. Our Hi-C saliency maps differ from Hi-C maps in that saliency maps tell us which contacts are *important*.

**Fig. 10:**
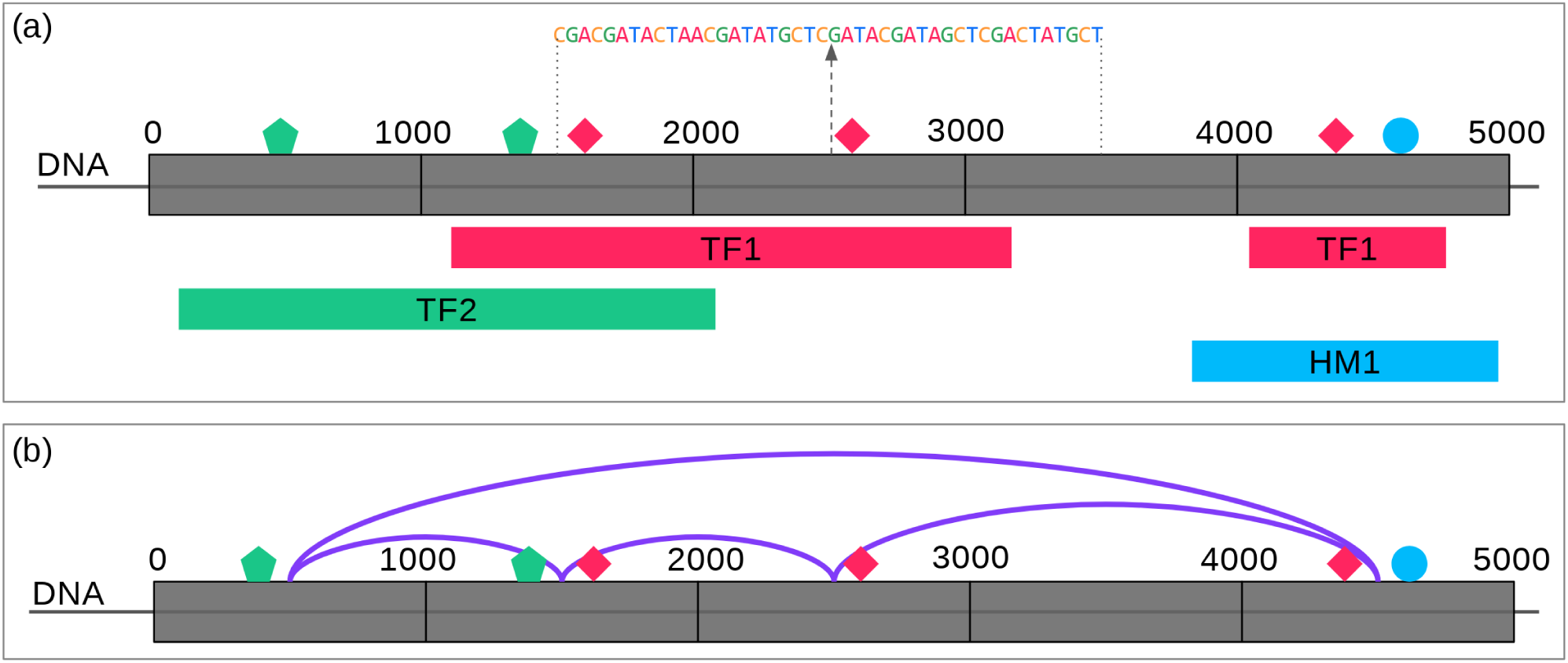
**(a) Sequence Data Processing** We extract 2000bp sequences surround 1000bp windows for any window that has an overlapping ChIP-seq peak. **(b) 3D Genome Data Processing** we use Hi-C contacts between the 1000bp windows from [23] as edges in our 3D genome graph.

## Footnotes

1 We use the terms epigenetic state and chromatin state interchangeably.

2 In our experiments, we use a different adjacency matrix **A** for each chromosome (intra-chromosome Hi-C maps). However, we generalize a **A** to represent all possible window interactions (i.e., including inter-chromosome maps).

